# Electrostatic repulsion causes anticooperative DNA binding between tumor suppressor ETS transcription factors and JUN-FOS at composite DNA sites

**DOI:** 10.1101/318048

**Authors:** Bethany J. Madison, Kathleen A. Clark, Niraja Bhachech, Peter C. Hollenhorst, Barbara J. Graves, Simon L. Currie

**Affiliations:** Department of Oncological Sciences; Huntsman Cancer Institute, University of Utah School of Medicine, Salt Lake City UT, 84112; Medical Sciences, Indiana University School of Medicine, Bloomington IN, 47405; Howard Hughes Medical Institute, Chevy Chase MD, 20815

**Author notes:** Both authors contributed equally to this work (Bethany J. Madison, Kathleen A. Clark). Present address: Department of Biophysics, University of Texas Southwestern Medical Center, Dallas, Texas 75390 (Simon L. Currie). To whom correspondence should be addressed: Barbara J. Graves.

**Keywords:** AP1 transcription factor (AP-1), ETS transcription factor family, gene expression, prostate cancer, protein-DNA interaction, protein-protein interaction, tumor suppressor genes

## Abstract

Many transcription factors regulate gene expression in a combinatorial fashion often by binding in close proximity on composite cis-regulatory DNA elements. Here we investigate the molecular basis by which ETS transcription factors bind with AP1 transcription factors JUN-FOS at composite DNA-binding sites. The ability to bind to DNA with JUN-FOS correlates with the phenotype of these proteins in prostate cancer: the oncogenic ERG and ETV1/4/5 subfamilies co-occupy ETS-AP1 sites with JUN-FOS *in vitro*, whereas JUN-FOS robustly inhibits DNA binding by the tumor suppressors EHF and SPDEF. EHF binds to ETS-AP1 DNA with tighter affinity than ERG in the absence of JUN-FOS, which may enable EHF to compete with ERG and JUN-FOS for binding to ETS-AP1 sites. Genome-wide mapping of EHF and ERG binding sites in a prostate epithelial cell line reveal that EHF is preferentially excluded from closely spaced ETS-AP1 DNA sequences. Structural modeling and mutational analyses indicate that adjacent positively-charged surfaces from EHF and JUN-FOS disfavor simultaneous DNA binding due to electrostatic repulsion. The conservation of positively charged residues on the JUN-FOS interface identified ELF1 as an additional ETS factor that exhibits anticooperative DNA binding, and we present evidence that ELF1 is frequently downregulated in prostate cancer. In summary, the divergence of electrostatic features of ETS factors at their JUN-FOS interface enables distinct binding events at ETS-AP1 DNA sequences. We propose that this mechanism can drive unique targeting of ETS transcription factors, thereby facilitating distinct transcriptional programs.

---

Sequence-specific transcription factors bind to cis-regulatory elements in enhancers and promoters to regulate gene expression. Composite DNA sequences consisting of multiple transcription factor binding sites enable precise and combinatorial control of gene transcription by integrating multiple inputs into a single transcriptional output (1,2). Multiple transcription factors can bind to a composite DNA sequence in a cooperative (tighter affinity for DNA), noncooperative (same affinity for DNA), or anticooperative (reduced affinity for DNA) manner. In many cases, transcription factors modulate the binding of each other at composite binding sites through protein-protein interactions (3–6), and/or through DNA-mediated effects(7–10). Although combinatorial regulation of gene transcription occurs frequently (4), the molecular basis of interplay between most transcription factor pairings at composite sites is poorly understood.

Closely-apposed sites for the binding of ETS and AP1 transcription factors play an important role in regulating cellular migration. These composite sites are found in the enhancers and promoters of genes such as the urokinase plasminogen activator (*PLAU*), the uridine phosphorylase (*UPP*), and the matrix metalloproteases (*MMP1*, *MMP9*, etc.)(11–14). Overexpression of “oncogenic” ETS factors from the ERG and ETV1/4/5 subfamilies occurs frequently in prostate cancers (15,16) and leads to the hyperactivation of ETS-AP1 regulated genes, ultimately resulting in enhanced cellular migration (11). Correspondingly, these oncogenic ETS factors bind to composite ETS-AP1 sites with JUN-FOS *in vitro* (17). Conversely, other ETS factors such as EHF and SPDEF function as tumor suppressors in prostate cancer (18–20) and repress the transcription of ETS-AP1 regulated genes (11,13). A simple hypothesis is that the tumor suppressor class of factors competes with oncogenic ones for binding to ETS-AP1 composite sites. However, no direct evidence for this hypothesis is available.

Here we investigate the difference between oncogenic and tumor suppressor ETS factors in binding to ETS-AP1 sites with JUN-FOS. Oncogenic proteins bound to composite sites with JUN-FOS in either a cooperative (ERG, FLI1) or non-cooperative manner (ETV1, ETV4). In contrast, the tumor suppressors EHF, SPDEF, and ELF1 displayed a robust anticooperative binding to DNA with JUN-FOS. In the absence of JUN-FOS, EHF bound to ETS-AP1 sequences with higher affinity than ERG suggesting that the inability of EHF to co-occupy DNA with JUN-FOS is not due to intrinsic DNA-binding differences. Genome-wide mapping of EHF and ERG DNA-binding sites in a prostate epithelial cell line provided support for anticooperative DNA binding between EHF and JUN-FOS at closely spaced ETS-AP1 composite sites. Structural modeling suggested that simultaneous DNA-binding would result in electrostatic repulsion between positive surfaces of EHF and JUN-FOS. In contrast, the corresponding surface on ERG is polar and negative, complementing the positive interface of JUN-FOS. In support of this model, mutation of the lysine residues in EHF enabled binding to DNA with JUN-FOS. Our results indicate that electrostatic properties regulate the ability of ETS factors to bind to composite ETS-AP1 DNA sequences with JUN-FOS, and implicate the divergence of these properties in the phenotypically diverse roles of ETS factors in prostate cancer.

## Results

### JUN-FOS differentially impacts ETS factor binding to composite ETS-AP1 sites

Oncogenic ETS factors, such as those from the ERG and ETV1/4/5 subfamilies, enhance transcription at ETS-AP1 regulated genes; conversely, ETS tumor suppressors, such as EHF and SPDEF, repress transcription from ETS-AP1 regulated genes (11,13,18,20). We hypothesized that differences in binding to DNA with JUN-FOS may be one reason for differential regulation by ETS factors. To test this hypothesis we expressed and purified full-length recombinant proteins for AP1 factors JUN and FOS, and for the ETS factors ETV1, ETV4, ERG, FLI1, EHF, and SPDEF. We measured the equilibrium dissociation constant (*K_D_*) for ETS proteins binding to an ETS-AP1 composite DNA sequence from the *UPP* promoter using electrophoretic mobility shift assays (EMSAs) in the presence and absence of JUN-FOS (Fig. 1A-C, Fig. S1, and Table S1). JUN-FOS enhanced the DNA binding of ERG and FLI1 (five to twenty-five fold), had minimal effects on the DNA binding of ETV1 and ETV4 (~ two fold), and strongly antagonized the DNA binding of EHF and SPDEF (greater than twenty fold). Therefore, different subgroups of ETS factors demonstrated cooperative, noncooperative, and anticooperative binding with JUN-FOS to an ETS-AP1 composite DNA sequence.

**Figure 1.**
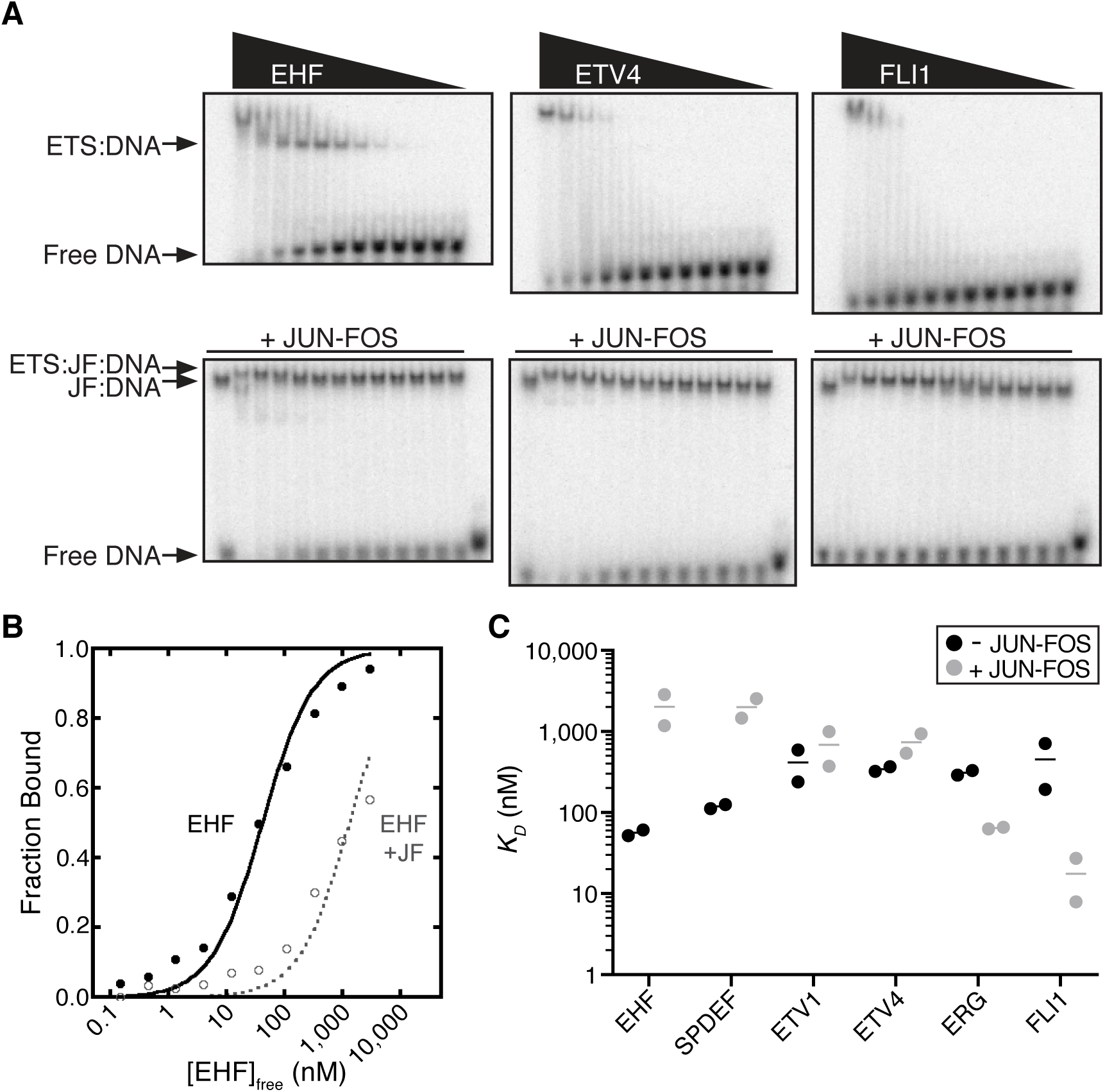
JUN-FOS differentially influences the DNA binding of ETS factors to AP1-ETS composite sites. *A,* Representative phosphorimages of electrophoretic mobility shift assays (EMSAs) for EHF (left), ETV4 (middle), and FLI1 (right) binding to the *UPP* promoter DNA duplex. ETS titrations were performed with DNA alone (top), and with nearly saturating JUN-FOS bound to the DNA (bottom). The higher band for EHF corresponds to two EHF molecules bound to the DNA duplex, as previously observed for similar ETS factors (17). *B,* Binding isotherms for EHF binding to *UPP* DNA in absence (black), and presence (gray) of JUN-FOS. Each data point is the mean from two experiments. *C, K_D_* values for ETS factors binding to *UPP* DNA without (black) or with (gray) JUN-FOS. The bar indicates the mean of the two experiments (filled circles). See Fig. S1 and Table S1 for quantification of equilibrium dissociation constants (*K_D_*) from EMSAs.

We also observed that ETS factors in the absence of JUN-FOS have distinct binding affinities for the ETS-AP1 sequence in the *UPP* promoter; EHF and SPDEF bound to this promoter with roughly ten-fold higher affinity than ERG, FLI1, ETV1, and ETV4 (Fig. 1C). We next tested ERG and EHF with a series of DNA sequences to further investigate this difference in binding to DNA. We found that ERG and EHF bind to a consensus high-affinity ETS binding sequence “SC1” (21,22) with similar affinities (Fig. 2A-C and Table S2). EHF again had an approximately ten-fold higher affinity for another ETS-AP1 DNA-binding sequence from the enhancer of *COPS8*. The proximal 5’ nucleotide outside of the core ETS motif is a cytosine (<u>C</u>GGAA) in the consensus ETS sequence, but is an adenosine (<u>A</u>GGAA) in many ETS-AP1 binding sites including those at the *UPP* promoter and *COPS8* enhancer (11,17). The nucleotide at this position has previously been shown to selectively affect the DNA binding of different ETS factors (22,23). To test whether this single nucleotide difference is important in selectively weakening ERG binding relative to EHF binding, we changed this nucleotide from cytosine to adenosine in the context of the high-affinity ETS sequence “SC1” (Fig. 2A)(21). This single change largely recapitulated the difference in binding affinities observed for the ETS-AP1 sequence. That is, ERG bound to this DNA with roughly ten-fold weaker affinity; whereas, the disruption of EHF binding was more subtle (Fig. 2B,C). Therefore, in the absence of JUN-FOS, EHF has a higher affinity for ETS-AP1 DNA sequences compared to oncogenic ETS factors. The tighter affinity of EHF may allow it to compete with ERG for ETS-AP1 DNA sequences despite binding anticooperatively with JUN-FOS.

**Figure 2.**
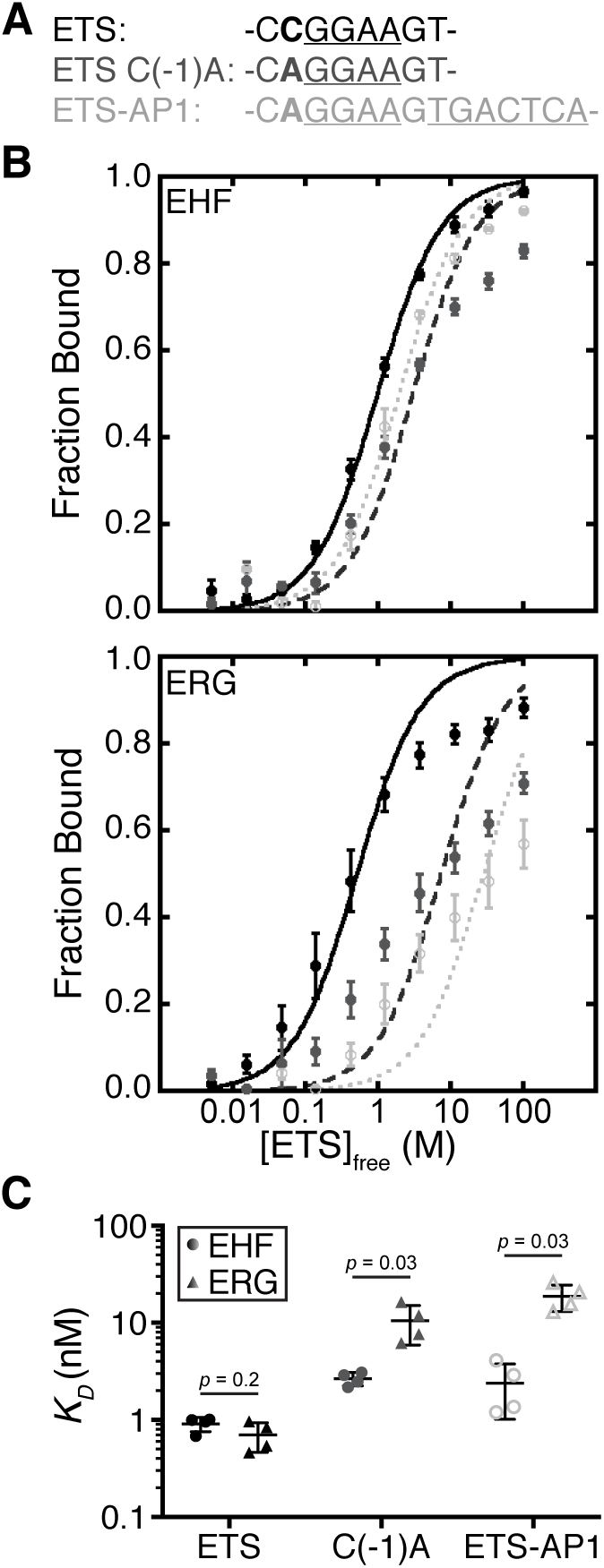
Single-nucleotide change flanking core ETS binding sequence differentially affects the DNA binding of ETS factors. *A,* DNA sequences used for measuring *K_D_* values with EHF or ERG. ETS and AP1 DNA-binding sites are underlined. *B*, Binding isotherms for EHF (top) and ERG (bottom) with the DNA duplex containing a consensus ETS DNA sequence for these factors (black), a single nucleotide present in ETS-AP1 composite motifs (dark gray), and an ETS-AP1 composite DNA sequence (light gray). Mean and standard deviation are displayed from at least four replicates. *C*, Comparison of *K_D_* values for EHF and ERG for all three DNA sequences. Each dot depicts a single experiment and lines depict the mean and standard deviation; see Table S2 for further quantification of EMSAs.

### ERG and EHF display differential preference for composite ETS-AP1 sites in vivo

To explore the biological significance of the cooperativity and anticooperativity displayed by ERG and EHF with JUN-FOS, respectively, we examined binding site preferences for these two proteins *in vivo*. Specifically, full-length ERG or EHF coding sequence was tagged with FLAG and expressed retrovirally in RWPE1 cells, a normal prostate epithelial cell line, and then FLAG-ChIPs were performed to determine chromatin occupancy genome-wide. ERG and EHF proteins were expressed at similar levels as judged by western blot analysis (Fig. S2A). Cluster analysis of the ChIP-seq data sets for FLAG-ERG and FLAG-EHF revealed four distinct groups - 1) regions with both high ERG and EHF occupancy; 2) regions with high EHF occupancy and low ERG occupancy; 3) regions with high ERG occupancy and low EHF occupancy; and 4) regions with low but significant occupancy for both proteins (Fig. 3A). Thus, ERG and EHF exhibit both redundant and unique genomic targets in RWPE1 cells.

**Figure 3.**
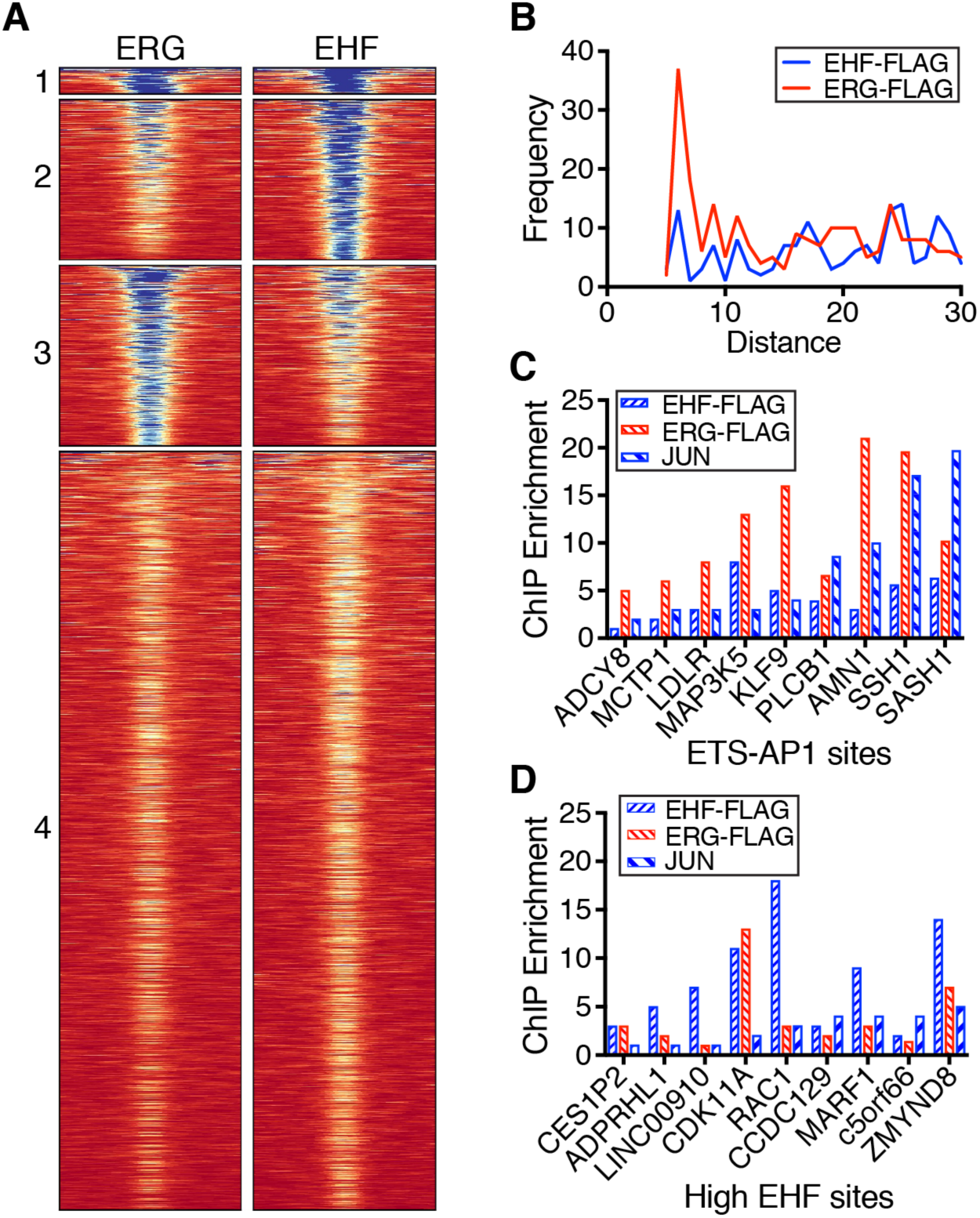
Preferential binding of ERG to ETS-AP1 sites *in vivo*. *A,* Heatmap of reads for ERG-FLAG and EHF-FLAG ChIP data; numbers to the left indicate clusters referred to in text. Analysis of ChIPseq data using MACS2 returned 34,746 enriched regions for ERG-FLAG and 44,977 for EHF-FLAG. *B,* Spacing between ETS and AP1 sites in top 1000 EHF-FLAG and ERG-FLAG ChIP peaks; spacing is defined as nucleotide distance between underlined residues in string below graph. Core nucleotides for ETS and AP1 sites are capitalized in string. *C,* qPCR quantification of EHF-FLAG, ERG-FLAG and JUN enrichment at putative regulatory elements for genes shown; regions selected based on match to ETS-AP1 sites with +6 spacing as shown in *B*. Two to three independent biological replicates provided similar patterns, but different maximum levels of enrichment. A representative experiment is shown. *D,* qPCR quantification of EHF-FLAG, ERG-FLAG and JUN enrichment at regions predicted to have high EHF occupancy based on ChIPseq data. ChIP enrichment for *C* and *D* is defined as the qPCR signal for that site divided by the qPCR signal for a neutral region, the 3’ UTR of *BCLxL1*.

Due to the high number of occupied regions, we limited further analysis to the top 1000 enriched regions for both proteins. Both ChIP datasets were enriched for DNA motifs matching the ETS binding consensus sequence (Fig. S2B), with slight differences in nucleotide preference surrounding the core GGA. The previously described composite “ETS-AP1 half-site” [CAGGAA(A/G)TGA] (11) was specifically enriched in the ERG dataset. Full AP1 sites (TGANTCA) were also overrepresented in both datasets (Fig. S2B). The composite element displaying tight spacing between ETS and AP1 motifs, which we examined by EMSAs, was more enriched in the ERG dataset as compared to the EHF dataset (Fig. 3B). Conversely, ETS-AP1 composite sites with more distant spacing were similarly represented in both ERG and EHF datasets. We conclude that anticooperative DNA binding between EHF and JUN-FOS occurs at ETS-AP1 composite sites with tight spacing.

To interrogate our genome-wide findings by quantifying differences in occupancy between ERG and EHF we performed direct ChIP-qPCR on randomly selected regions from the ERG-FLAG ChIP data that had an ETS-AP1 site with tight spacing. JUN occupancy was also confirmed in these direct ChIPs. Occupancy was defined by ChIP enrichment, the ratio of the PCR signal from a test region compared to a control region. At seven of the eight ETS-AP1 sites assayed, ChIP enrichment was at least two-fold and up to seven-fold higher for ERG than EHF (Figs. 3C, S2C); thus, ERG-FLAG occupancy at regions with composite ETS-AP1 sites was higher than FLAG-EHF occupancy, as suggested by the motif analysis in the genome-wide analysis. JUN occupancy was similar at most ETS-AP1 sites in RWPE1 cells expressing either ERG or EHF suggesting that the presence of JUN-FOS may be deterring EHF binding (Fig. S2D). Additionally, JUN was depleted from two ETS-AP1 sites at enhancers for *SASH1* and *PLCB1* in EHF expressing cells indicating that EHF may prevent JUN-FOS from binding to select ETS-AP1 sites. In order to verify that lower EHF occupancy at ETS-AP1 sites did not reflect a detection problem of EHF versus ERG in cells, we assayed regions that were specifically enriched for EHF in the cluster analysis. At specific EHF-enriched sites EHF occupancy was equal or greater than ERG (Fig. 3D). JUN occupancy was also lower at these regions compared to regions with preferential ERG binding. Collectively, these data suggest that ERG and EHF have distinct DNA-binding profiles in prostate cancer cells, including the relative depletion of EHF at closely spaced ETS-AP1 composite sites.

### Positive residues N-terminal of, and within, the ETS domain of EHF mediate anticooperative binding to DNA with JUN-FOS

To characterize the anticooperative binding to DNA with JUN-FOS, which we had observed for both EHF and SPDEF, we chose EHF for mapping the minimal regions of EHF and JUN-FOS that were sufficient for anticooperative DNA binding. The DNA-binding domains of JUN (JUN^ΔN250^ ^ΔC319^) and FOS (FOS^ΔN131^ ^ΔC203^) were sufficient for antagonizing the DNA binding of full-length EHF (Fig. S3). Residues 193-300 of EHF (EHF^ΔN193^), which includes the ETS domain and 16 residues N-terminal of the ETS domain, retained the full anticooperative binding behavior of the full-length protein (Figure 4A-B). Removal of the flanking N-terminal residues (EHF^ΔN203^) resulted in approximately ten-fold stronger binding to DNA in the presence of JUN-FOS, although the minimal ETS domain still retained five-fold anticooperative behavior. Therefore, the minimal regions required for anticooperative DNA binding for JUN and FOS are the DNA-binding domains; whereas in the case of EHF, both the ETS domain and a proximal N-terminal region are needed.

**Figure 4.**
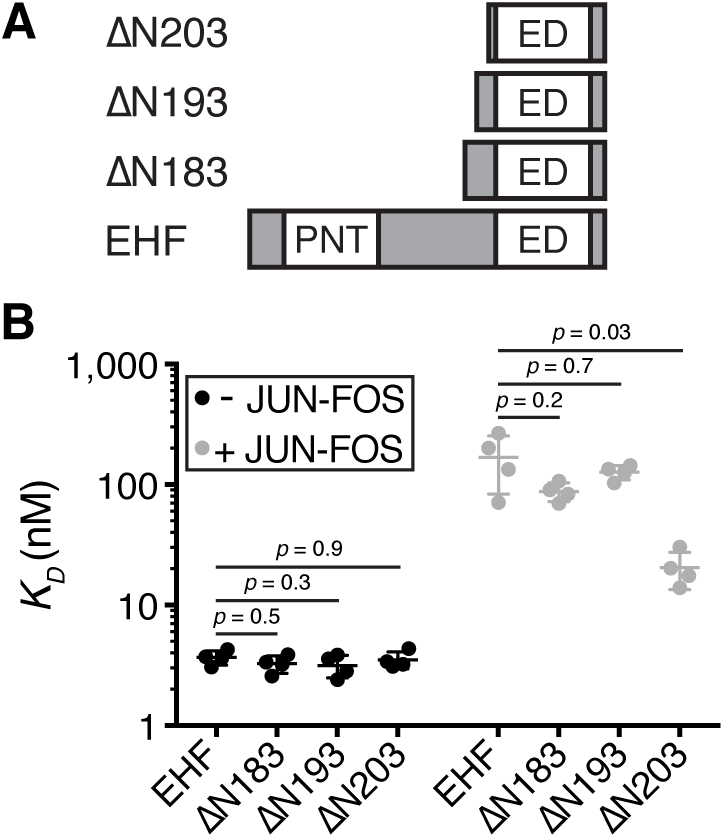
Sequences N-terminal of the ETS domain and within the ETS domain of EHF are important for anticooperative binding with JUN-FOS to composite ETS-AP1 sites. *A*, Schematic of EHF truncation series. ETS DNA-binding domain (ED) and Pointed domain (PNT) are labeled. *B*, *K_D_* values for EHF and N-terminal truncations binding to an ETS-AP1 sequence without (black) and with (gray) JUN-FOS. The mean and standard deviation are represented by horizontal lines; see Table S5 for further quantification of EMSAs

We next generated a structural model to interrogate the differential binding between ETS factors with JUN-FOS at composite ETS-AP1 sites using previously characterized DNA-bound structures of JUN-FOS (24) and ETS factors ERG (25) and ELF3 (26), which is a close homolog of EHF. In order to model the ternary complex we aligned the DNA sequences to mimic a common composite site with the ETS site just upstream of the AP1 site (GA<u>GGAA</u>G<u>TGACTCA</u>)(11). This modeling demonstrated no significant steric overlap for the ETS domain of EHF or ERG binding with JUN-FOS to the composite motif. However, the regions on the ETS domains of EHF and ERG in closest proximity to JUN-FOS have contrasting charge properties. For EHF, the N-terminus of the ETS domain, the loop between α-helices H2 and H3, and the C-terminus of H3 are all positively charged; whereas, the analogous regions in ERG are neutral or negatively charged (Fig. 5A,B and Fig. S4). The regions of JUN and FOS proximal to ETS factors are positively charged. Therefore, our modeling suggests that the positively-charged interfaces of EHF and JUN-FOS would cause electrostatic repulsion, disfavoring simultaneous binding to composite DNA sequences. In contrast, the lack of positive charges in the ERG interface presents a more favorable interaction surface for JUN-FOS allowing for concurrent binding with JUN-FOS to composite DNA sequences.

**Figure 5.**
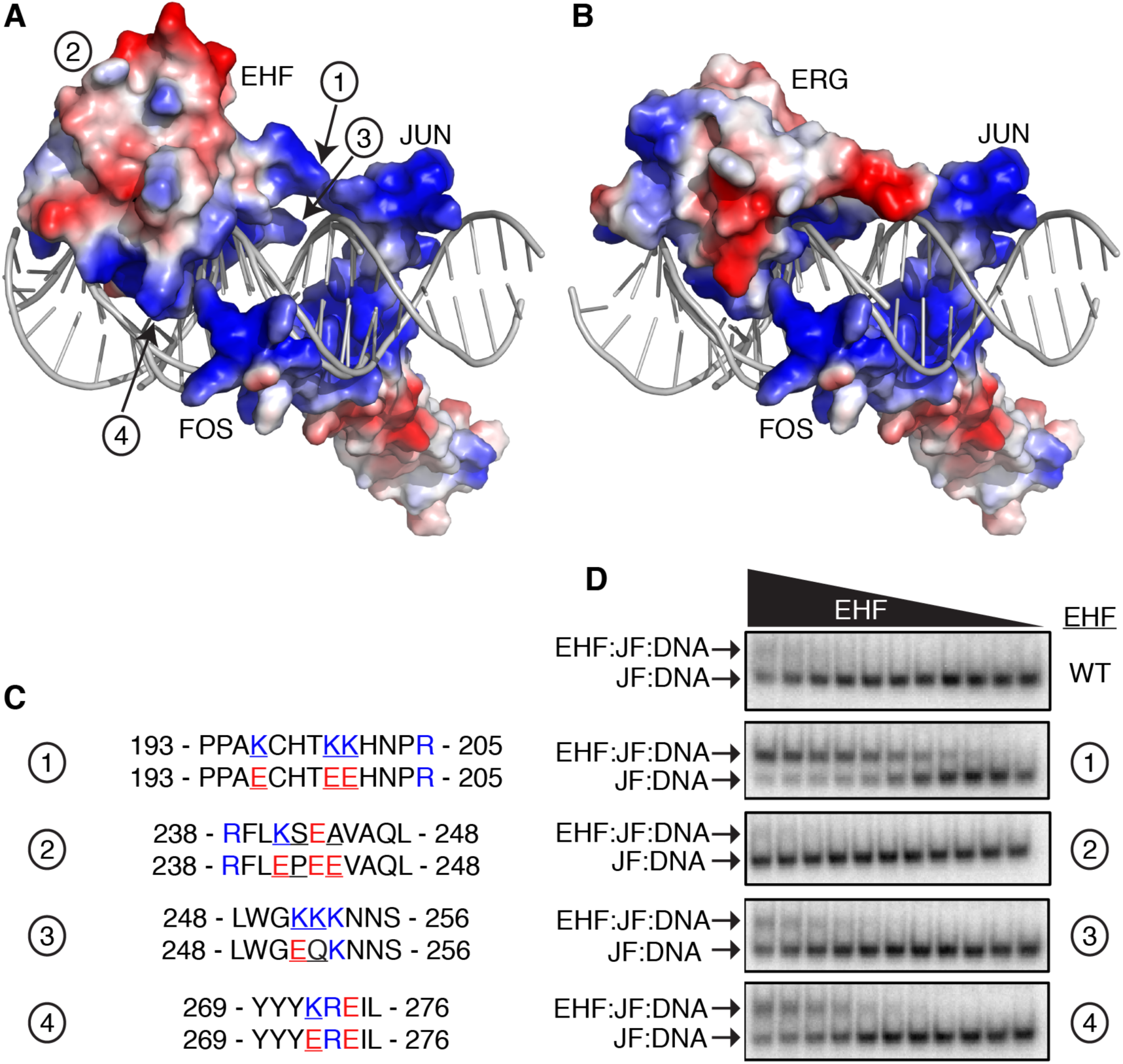
Positively-charged residues near the JUN-FOS interface are important for anticooperative binding of EHF and JUN-FOS. Structural model of EHF *A,* and ERG *B*, binding to an ETS-AP1 composite DNA sequence with JUN-FOS. EHF, ERG, and JUN-FOS are shown in surface mode and colored according to electrostatic potential (red, negative; blue, positive). Regions of EHF that were subsequently mutated are labeled 1 – 4. Note that EHF residues 193-204 are not present in this structure. *C*, Listing of EHF residues that were mutated. Circled numbers 1 – 4 correspond to the regions labeled in *A*. The top and bottom sequences indicate the native and mutated residues, respectively. Residues are colored according to charge, as in *A*. *D*, Portions of EMSAs showing EHF wildtype (WT) and mutants bound in the presence of JUN-FOS on an ETS-AP1 site. EHF was serially diluted in twofold increments. Bands corresponding to JUN-FOS bound to DNA (JF:DNA) as well as EHF and JUN-FOS bound to DNA (EHF:JF:DNA) are labeled. See Fig. S6 and Table S6 for further quantification of EMSAs.

Next we tested the functional importance of positive residues within EHF for anticooperative binding with JUN-FOS to composite sites. Four positively-charged regions of EHF were mutated: the N-terminal region preceding the ETS domain (Lys196Glu; Lys200Glu; Lys201Glu), the loop between β-strand S2 and α-helix H2 (Lys241Glu; Ser242Pro; Ala244Glu), the loop between α-helices H2 and H3 (Lys251Glu; Lys252Gln), and the C-terminal end of α-helix H3 (Lys272Glu) (Fig. 5C). These regions were selected based on being more positively charged in ETS factors that displayed anticooperative binding with JUN-FOS (Fig. S4 and Fig. S5). The EHF residues within the ETS domain were mutated to corresponding residues in ETS factors that bind with JUN-FOS to composite sites, and the lysine residues in N-terminal region preceding the ETS domain were mutated to glutamate residues, as there is no meaningful sequence alignment for this region between different subfamilies of ETS factors (Fig. S5). Mutant proteins were tested for binding to DNA alone, and with saturating amounts of JUN-FOS. Importantly, these mutations did not significantly change the binding of EHF to DNA in the absence of JUN-FOS (Fig. S6). However, mutating the region N-terminal of the ETS domain completely ablated anticooperative DNA binding with JUN-FOS (Fig. 5D and Fig. S6). Mutation of H3, and to a lesser extent the H2-H3 loop, showed lower impact. As a control, mutation of the S2-H2 loop, which is on the opposite side of the ETS domain from JUN-FOS in this ETS-AP1 composite motif arrangement, did not alter EHF binding with JUN-FOS. Therefore, positive residues in EHF that form the JUN-FOS interface are important for the anticooperative binding of EHF and JUN-FOS to ETS-AP1 DNA sequences.

### Charge-based prediction of additional ETS tumor suppressors

Based on the importance of positive residues for the anticooperative binding of EHF to DNA with JUN-FOS, we attempted to predict based on protein sequence which other ETS factors could display similar binding behavior. We selected ERF, GABPA, ELF1, and ELK4 for further analysis as these factors represent ETS factor subfamilies that have not been examined for binding with JUN-FOS previously (17) or in this study (Fig. S7). Like EHF, ELF1 has positive residues in all of the three positions that contribute to anticooperative binding with JUN-FOS (Fig. S5). In contrast, ERF, GABPA, and ELK4 lack positive residues in at least one of these important regions. JUN-FOS antagonized ELF1 DNA binding, enhanced ERF binding to DNA, and had little effect on GABPA and ELK4 (Fig. 6A). These additional data allowed us to predict the effect of JUN-FOS on the remaining untested ETS factors based on sequence homology (Figs. S5, S7). Comparing the charge of ETS domains and flanking regions with DNA binding with JUN-FOS demonstrates that ETS factors that anticooperatively bind to DNA with JUN-FOS tend to be more positively charged than other ETS factors (Fig. 6B). These data suggest that ELF1 binds anticooperatively with JUN-FOS due to a similar electrostatic repulsion mechanism as observed for EHF.

**Figure 6.**
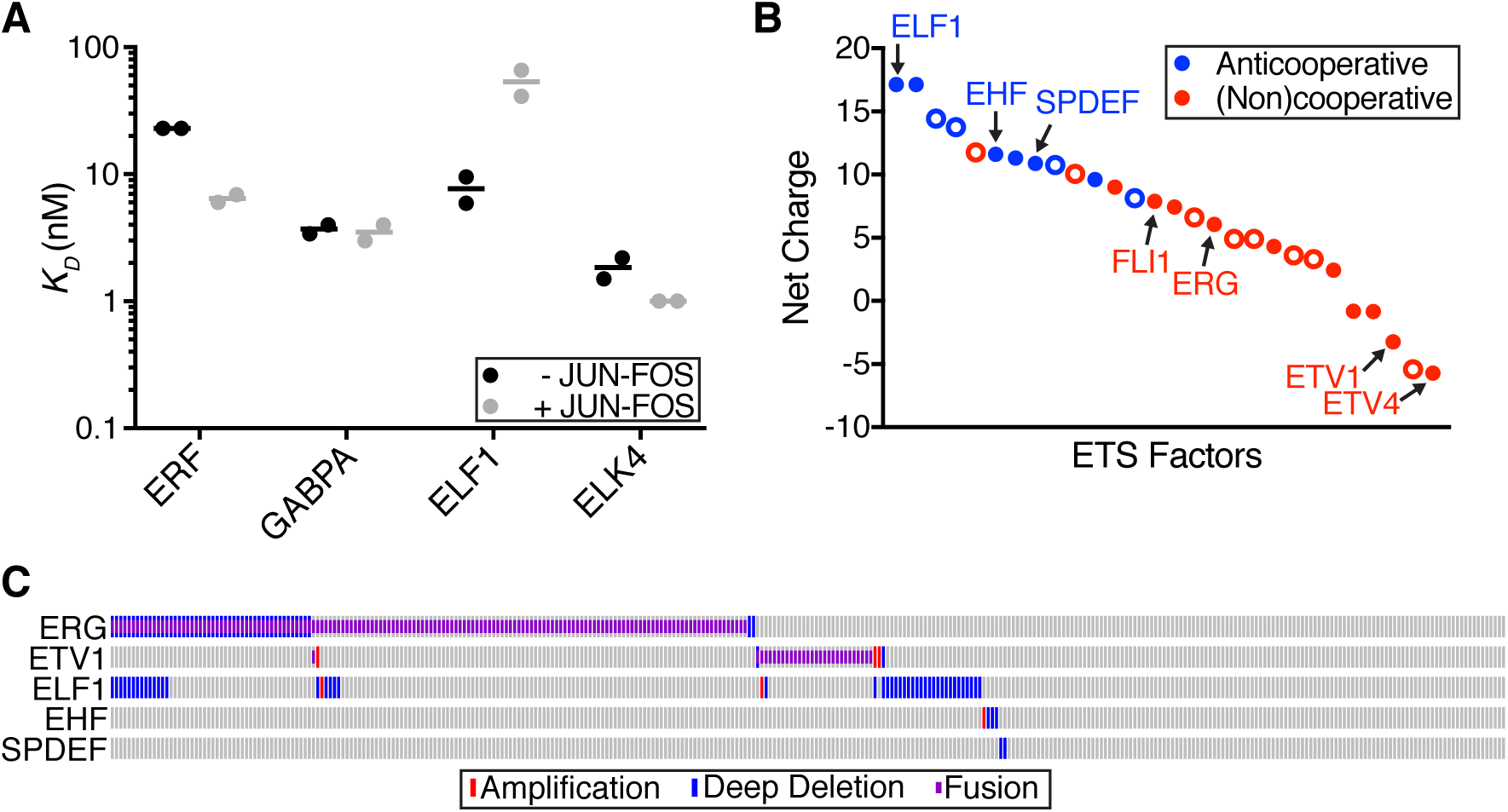
Prediction of novel ETS tumor suppressors in prostate cancer based on protein charge. *A*, Comparison of *K_D_* values for ERF, GABPA, ELF1, and ELK4 alone (black) and with near saturating amounts of JUN-FOS (gray). Filled circles represent an individual experiment and lines represent the mean. See Table S7 for *K_D_* values. *B,* Plot of ETS factors indicating the charge of the ETS domain and flanking sequences, and the DNA binding behavior with JUN-FOS. Filled circles represent experimental evidence from this study and from Kim et al., 2007; open circles represent predictions based on sequence homology (Fig. S5). Blue indicates anticooperative binding and red indicates non-cooperative or cooperative binding. *C,* Example of an oncoprint curated from cBioPortal showing mutational frequencies of the ETS factors ERG, ETV1, ELF1, EHF, and SPDEF (http://www.cbioportal.org)(29,30). This particular example is from a TCGA prostate cancer study published in 2015 (32). ERG and ETV1 are frequently overexpressed through gene fusions and EHF and SPDEF are rarely present in deep deletions, as previously characterized (15,16,19,20). Interestingly, ELF1 is also frequently involved in deep deletions suggesting that it may be a tumor suppressor in prostate cancer. See Figure S8 for additional studies that found frequent ELF1 gene deletions in prostate cancer patients.

We originally examined the anticooperative ETS factors EHF and SPDEF based on their reported tumor suppressor roles in prostate cancer (18,20). Thus, we hypothesized that other positively charged ETS factors that bind to DNA in an anticooperative manner with JUN-FOS might also behave as tumor suppressors in prostate cancer. In support of this hypothesis, protein levels of ELF and SPI subfamily members ELF1, ELF2, ELF4, and SPIB are reported to be often downregulated in prostate cancer samples (Fig. S8A) (27,28). At the genomic level, *ELF1* in particular is affected by deep gene deletions in up to 20% of prostate cancer samples from different prostate cancer studies (Figs. 6C and S8B)(29–36). Cumulatively, these data suggest that positively charged ETS factors that bind to consensus DNA sites with JUN-FOS in an anticooperative manner may behave as tumor suppressors in prostate cancer.

## Discussion

Here we report the variable binding of ETS transcription factors with JUN-FOS to composite DNA sequences. DNA binding of the tumor suppressors EHF, ELF1, and SPDEF is strongly antagonized by JUN-FOS, in contrast to oncogenic factors from the ERG and ETV1/4/5 subfamilies. We propose that this difference in binding to DNA with JUN-FOS contributes to the opposing regulation of ETS-AP1 regulated genes with roles in cellular migration by oncogenic and tumor suppressor ETS factors (Fig. 7A)(11,13).

**Figure 7.**
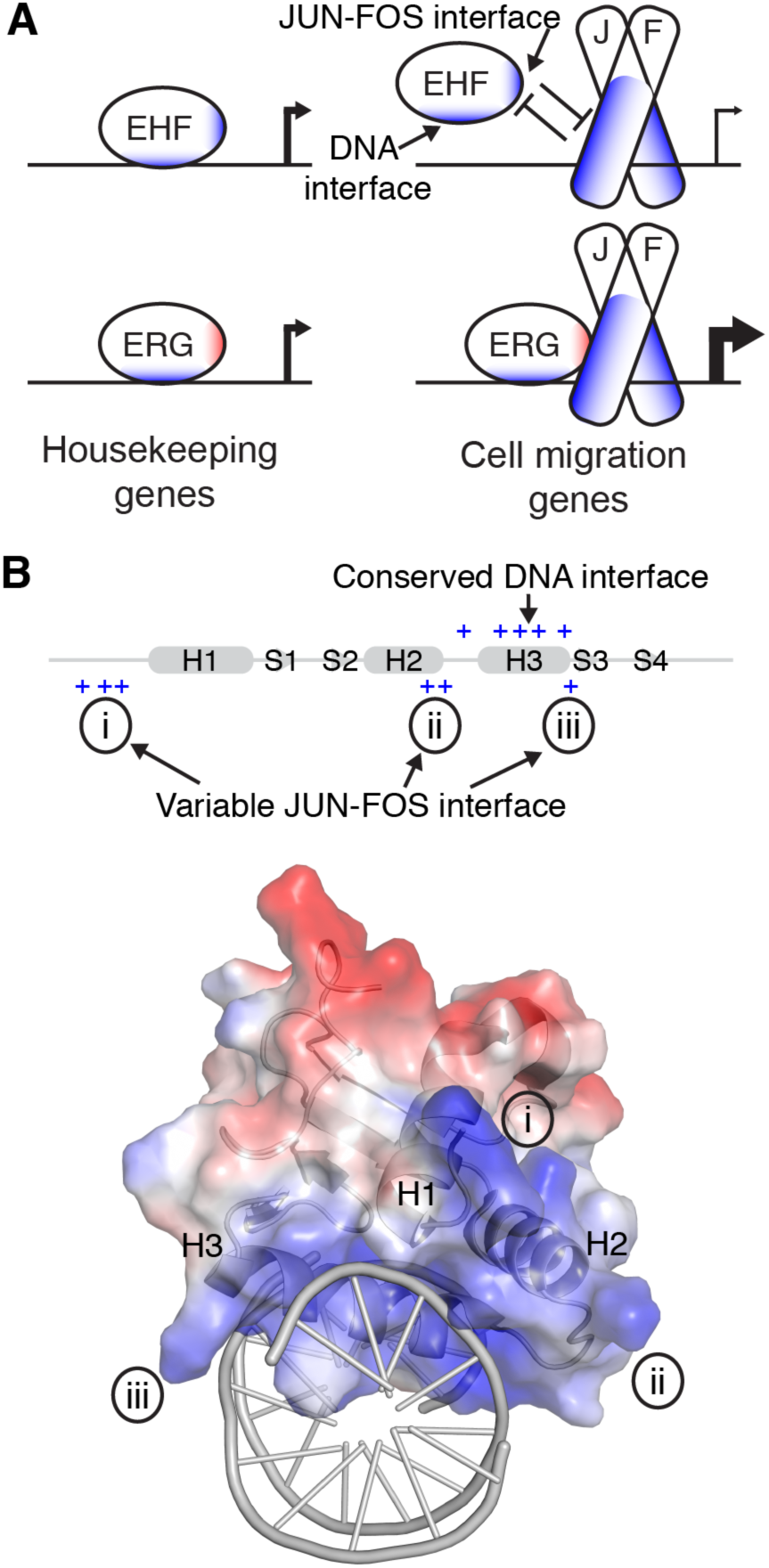
*A*, Model for differential regulation of ETS-AP1 sites by ETS factors. Left, EHF and ERG have DNA-binding surfaces similar to all ETS factors, and therefore bind to ETS DNA sequences with relatively similar affinities. Right, the distinct JUN-FOS interface of EHF and ERG allows ERG to bind to ETS-AP1 sequences with JUN-FOS, but prevents EHF from binding to ETS-AP1 sequences with JUN-FOS. This difference in binding affinities is consistent with the repression and activation of ETS-AP1 regulated genes by EHF and ERG, respectively (11,13). *B*, Three positive regions of EHF form the JUN-FOS interface. Top, the ETS domain of EHF is depicted in cartoon format; cylinders and arrows indicate α-helices and β-strands, respectively, and are named according to previous nomenclature (37). Positive residues in α-helix H3 are at the primary DNA interface and are highly conserved amongst human ETS factors (Figs. S5 and S7). In contrast, positive residues N-terminal of the ETS domain (i), in the H2-H3 loop (ii), and C-terminal of α-helix H3 (iii) form the JUN-FOS interface and are only found in a subset of human ETS factors. Bottom, these three regions of EHF, which are separated in primary sequence, converge to form a broad positively-charged interface for JUN-FOS. See Fig. S9 for the JUN-FOS interfaces of SPDEF, ELF1, and ERG.

### Model for anticooperative DNA binding between EHF and JUN-FOS

Residues flanking and within the ETS domain of EHF contribute to anticooperative DNA-binding with JUN-FOS, as revealed by a truncation series. In particular, the region including multiple lysine residues just N-terminal of the ETS domain is the single strongest contributor to this effect. Structural modeling indicated that the N-terminus of the ETS domain as well as the loop between α-helices H2 and H3 and the C-terminal end of H3 are all positioned at the JUN-FOS interface when both factors are bound to DNA. These three regions of ETS factors, though separate in the primary sequence, converge in the tertiary structure forming a tripartite interface. In EHF, all of these regions are positively charged and are positioned near positively-charged regions of the DNA-binding domains of JUN and FOS. Therefore, we suggest that electrostatic repulsion between basic EHF and JUN-FOS surfaces inhibits simultaneous DNA binding in this orientation. In contrast, the collective JUN-FOS interface on ERG is composed of negative and polar residues, making ERG more suitable for simultaneous DNA binding with JUN-FOS. Mutation of EHF to eliminate basic residues along this interface at any of the three regions abrogated anticooperative binding with JUN-FOS. Eliminating basic residues from the region N-terminal of the ETS domain again demonstrated the largest effect, matching the truncation series data. However, mutation of the H2-H3 loop or the C-terminus of H3 also significantly improved the DNA binding of EHF with JUN-FOS, indicating that positive residues at all three sites contribute to the full anticooperative effect. Interestingly, further examination of additional ETS factors demonstrated that ELF1, which is positively charged at all three analogous regions, exhibited anticooperative DNA-binding with JUN-FOS. In contrast, ERF and ELK4, which lack positively-charged residues in one of these regions, do not exhibit anticooperative DNA binding. Lastly, introduction of the positive residues from EHF into any single site of the tripartite interface on ERG (i.e. N-terminal, H2-H3 loop, or C-terminus of H3) failed to transfer any level of anticooperative binding with JUN-FOS into ERG (data not shown). Therefore, we propose that anticooperative binding to DNA with JUN-FOS requires three different stretches of positive residues on EHF that cumulatively form a basic interface with JUN-FOS (Fig. 7B and Fig. S9).

Despite close proximity to the DNA interaction surface, elimination of basic residues from the JUN-FOS interface of EHF had no significant impact on DNA binding in the absence of JUN-FOS. We interpret this to indicate that DNA binding alone and anticooperative DNA binding with JUN-FOS are fundamentally different and separable properties. Correspondingly, the arginine and tyrosine residues along the core DNA-recognition surface of α-helix H3 are highly conserved across ETS factors (Figs. S5, S9)(22,37). Anticooperative ETS factors, such as EHF, ELF1, SPDEF, and SPI1, bind with slightly stronger affinities to ETS-AP1 DNA sequences in the absence of JUN-FOS compared to cooperative ETS factors. These data suggest that when present in excess anticooperative ETS factors will compete with oncogenic ETS factors and JUN-FOS for binding to ETS-AP1 sites. In principle, this competition would reduce the level of transcription from ETS-AP1 regulated genes by reducing the occupancy of oncogenic ETS factors and/or JUN-FOS (Fig. 7A). This hypothesis is supported by previously reported data indicating that anticooperative ETS factors are highly expressed in normal prostate cells and dampen the transcription of genes involved in cellular migration that are regulated by ETS-AP1 sites (11,13,17,38). Therefore, we propose that anticooperative DNA binding with JUN-FOS is one mechanism for tumor suppressor ETS factors to repress the transcription of ETS-AP1 regulated genes.

### Distinct mechanisms of transcriptional repression among ETS factors

A recent report implicated the ETS factor ERF as a novel tumor suppressor in prostate cancer and suggested that ERF represses transcription from ERG and androgen receptor (AR) target genes by competing with ERG for binding to ETS DNA sequences (39). Our analysis of an ERF truncation (residues 1-126) indicates that ERF cooperatively binds to ETS-AP1 DNA sequences with JUN-FOS, unlike the other ETS tumor suppressors that we tested. Although we cannot rule out that the full-length ERF may behave differently, several lines of evidence support that ERF transcriptional repression occurs through a distinct mechanism as compared to EHF, ELF1, and SPDEF. The repressor domain of ERF is distal from the ETS DNA binding domain, greater than 350 residues away in primary sequence, and is fully transferable to a heterologous protein (40). In contrast, the positive residues of EHF that facilitate anticooperative DNA binding with JUN-FOS, flank or are within the DNA-binding domain, and are not easily transferable to other proteins, such as ERG. Furthermore, ERF acts as a transcriptional repressor in all contexts tested thus far (40); whereas, EHF represses or activates genes in a context-dependent manner (13,41). These data suggest that ERF-mediated transcriptional repression occurs at a level other than DNA-binding, such as through the recruitment of corepressors for example. In contrast, our results suggest that anticooperative binding to DNA with JUN-FOS is a distinct mechanism of transcriptional repression that may contribute to the tumor suppressor phenotypes of the ETS factors EHF, SPDEF, and ELF1 in prostate cancer (18–20,33,42).

### Diversification of interaction surfaces and functional regulation

Individual ETS transcription factors display diverse developmental and disease-related phenotypes (37,43), yet possess ETS domains with remarkably similar DNA-binding preferences (22). How might this apparent contradiction be explained? One route for specificity involves the use of composite DNA sequences for combinatorial regulation. For example, composite ETS-RUNX sites are found in T-cell activation genes, and ETS1 specifically regulates these genes through cooperatively binding to DNA with RUNX1 (6,10,23,44). ERG and FLI1 bind cooperatively to DNA with JUN-FOS, although the molecular basis for this cooperativity remains unclear (17). Here we demonstrate that a subset of ETS factors, including SPDEF, EHF, and ELF1, bind to DNA in an anticooperative manner with JUN-FOS. This anticooperativity is dependent on positively charged residues from multiple regions of the protein that together form the JUN-FOS interface. The JUN-FOS interacting surface is distinct from the conserved DNA-binding surface, enabling precise control on modulating transcriptional activity at genes regulated by ETS-AP1 composite sites without impacting other genes that are regulated by ETS sites. An analogous electrostatic repulsion mechanism has recently been described for the selective recognition of appropriate substrates by tyrosine kinases involved in T-cell signaling (45,46). Therefore, electrostatic selection may be a general mechanism for fine-tuning molecular interactions that contributes to the phenotypic diversity of large gene families.

## Materials and Methods

### Expression plasmids

Open reading frames corresponding to full-length or truncated JUN, EHF, ELF1, ELK4, ERF, ERG, ETV1, ETV4, FLI1, FOS, GABPA, and SPDEF were cloned into the bacterial expression vector pET28 as previously described (47,48). Point mutations were introduced into the EHF ΔN193 plasmid using the QuikChange site-directed mutagenesis protocol (Stratagene).

### Expression and purification of proteins

ETS proteins were expressed in *Escherichia coli* (λDE3) cells. Full-length EHF and truncated ETS proteins (see Table S8 for protein sequences) were efficiently expressed into the soluble fraction. One-liter cultures of Luria broth (LB) were grown at 37°C to an OD600 of approximately 0.7. The cultures were then induced with 0.5 mM isopropyl-β-D-thiogalactopyranoside (IPTG) for approximately three hours at 30°C. Cultures were centrifuged at 12,000 x *g* for 10 min at 4°C. Cells were resuspended with 25 mL buffer (per L of culture) containing 25 mM Tris pH 7.9, 1 M NaCl, 0.1 mM ethylenediaminetetraacetic acid (EDTA), 2 mM 2-mercaptoethanol (βME), and 1 mM phenylmethanesulfonylfluoride (PMSF), flash frozen in liquid nitrogen, and stored at - 80°C. Cells were lysed by sonication and centrifuged at 125,000 x *g* for 30 min at 4°C. The supernatant containing the soluble fraction was loaded onto a Ni^2+^ affinity column (GE Biosciences) and eluted over a 5 to 500 mM imidazole gradient. Fractions containing purified protein were pooled and dialyzed overnight at 4°C into a buffer containing 25 mM Tris pH 7.9, 10% glycerol (v:v), 1 mM EDTA, 50 mM KCl, and 1 mM dithiothreitol (DTT). After centrifugation at 125,000 x *g* for 30 min at 4°C, the soluble fraction was loaded onto a SP-Sepharose cation exchange column or Q-Sepharose anion exchange column (GE Biosciences) depending on the isoelectric point of the individual protein, then eluted by a linear gradient from 50 mM to 1M NaCl. Fractions containing purified protein were pooled and further purified by size-exclusion chromatography on a Superdex 75 column run with a buffer containing 25 mM Tris pH 7.9, 10% glycerol, 1 mM EDTA, and 300 mM KCl. Fractions containing purified protein were pooled and concentrated by 30-kDa, 10-kDa, or 3-kDa molecular weight cut-off (MWCO) Centricon devices (Sartorius). Concentrated proteins were snap frozen with liquid nitrogen and stored at -80°C in single-use aliquots for subsequent EMSA studies.

Full-length ERG, FLI1, ETV1, ETV4, and SPDEF were predominantly expressed into inclusion bodies (See Table S8 for protein sequences). Protein expression was induced, then cells were centrifuged and stored as described above, with the exception of ETV4, which was induced by autoinduction as described previously (48,49). After the initial sonication, the samples were centrifuged at 31,000 x *g* for 15 min at 4°C, then the soluble fraction was discarded. This procedure was performed a total of three times to wash the inclusion bodies in 25 mM Tris pH 7.9, 1 M NaCl, 0.1 mM EDTA, 5 mM imidazole, 2 mM βME, and 1 mM PMSF. The final insoluble pellet was re-suspended in a buffer containing 25 mM Tris pH 7.9, 1 M NaCl, 0.1 mM EDTA, 5 mM imidazole, 2 mM βME, 1 mM PMSF, and 6 M urea using sonication. After rotation for ~ 1 hr at 4°C, the sample was centrifuged at 125,000 x *g* for 30 min at 4°C. The soluble fraction was loaded onto a Ni^2+^ affinity column and refolded on-column by switching to the same buffer lacking urea. After elution with a 5 to 500 mM imidazole gradient, the remaining purification steps (ion-exchange and size-exclusion chromatography) were performed, as described above.

Full-length JUN and FOS proteins were expressed and purified as described previously (50,51). Briefly, JUN and FOS expressed into the insoluble fraction and were expressed and purified as above, with the following exceptions. FOS was expressed in Rosetta 2 cells for supplementation of rare Arg tRNAs. Inclusion bodies were purified and solubilized as described above, then JUN and FOS were combined for JUN-FOS heterodimers, diluted to 200 ng/μL (total protein), then dialyzed for at least 3 hr each against the following three buffers (in sequential order): (1) 25 mM Tris pH 6.7, 0.1 mM EDTA, 10% glycerol, 5 mM βME, 1 M NaCl, 1 M urea; (2) same as (1) but without urea; (3) same as (2) but with NaCl reduced to 100 mM. Refolded samples were then purified by Ni^2+^ affinity and size-exclusion chromatography, as described above.

Truncated JUN^ΔN250 ΔC319^ and FOS^ΔN131^ ΔC203 proteins expressed into the insoluble fraction and solubilized, as described above. JUNΔN250 ΔC319 and FOSΔN131 ΔC203 were then individually loaded onto a Ni^2+^ affinity column, refolded on-column, and eluted, as described above. JUNΔN250 ΔC319 and FOSΔN131 ΔC203 were then combined to form heterodimers and further purified by size-exclusion chromatography, as described above.

### Electrophoretic mobility shift assays (EMSA)

DNA-binding assays of ETS factors utilized duplexed oligonucleotides corresponding to the promoter or enhancer regions of the genes *UPP1* and *COPS8*, respectively, and a consensus high-affinity ETS binding site SC1 (<u>S</u>elected <u>C</u>lone 1) (11,17,21). The DNA sequences for these oligonucleotides are listed in Table S8. Each pair of oligonucleotides, at 2 μM as measured by absorbance at 260 nm on a NanoDrop 1000 (Thermo Scientific), were labeled with [γ-^32^P] ATP using T4 polynucleotide kinase at 37°C for 30 min. After purification over a Bio-Spin 6 chromatography column (Bio-Rad), the oligonucleotides were incubated at 100°C for 5 min, and then cooled to room temperature over ~ 2 hr. The DNA concentration for EMSAs was diluted to 5 x 10^-11^ M and held constant. JUN-FOS was titrated against each DNA sequence to determine near saturating amounts, where bound DNA was approximately 80% of total DNA. For the *UPP* promoter, full-length JUN-FOS was included at 1 μM while the truncated JUN-FOS was included at 100 nM for the *COPS8* enhancer. ETS factor concentrations were titrated from the μM to sub-nM range in order to determine the equilibrium dissociation constant (*K_D_*) for ETS factors against DNA and JUN-FOS:DNA complexes. Protein concentrations were determined after thawing each aliquot of protein using the Protein Assay Dye Reagent (Bio-Rad). The binding reactions were incubated for 3 hr at 4°C in a buffer containing 25 mM Tris pH 7.9, 0.1 mM EDTA, 60 mM KCl, 6 mM MgCl2, 200 μg/mL Bovine Serum Albumin, 10 mM DTT, 100 ng/μL poly (dIdC), and 10% (v:v) glycerol. Reactions were resolved on a 4% or 6% (w:v) native polyacrylamide gel run at 4°C for experiments with full-length or truncated JUN-FOS, respectively. The ^32^P-labeled DNA was quantified on dried gels by phosphorimaging on a Typhoon Trio Variable Mode Imager (Amersham Biosciences). Equilibrium dissociation constants (*K_D_*) were determined by nonlinear least squares fitting of the total ETS protein concentration [P]_t_ versus the fraction of DNA bound ([PD]/[D]_t_) to the equation [PD]/[D]_t_ = 1/(1+*K_D_*/[P]_t_) with Kaleidagraph (v. 3.51; Synergy Software). Due to the low concentration of total DNA, [D]_t_, in all reactions, the total protein concentration is a valid approximation of the free, unbound protein concentration. *K_D_* values are reported as mean ± standard deviation. Statistical *t* tests were calculated with Graph Pad Prism (Version 7.0b) for experiments with at least three replicates. Consistent standard deviation was not assumed, and the two-stage step-up false discovery rate (FDR) approach of Benjamini, Krieger, and Yekutieli was used with a desired FDR of 1% (52).

### Cell culture and viral expression

RWPE1 cells were obtained from American Type Culture Collection and cultured accordingly. Full-length ERG and EHF cDNAs with an added C-terminal 3x FLAG tag were cloned into a modified pLHCX retrovial expression vector (Clontech) with the CMV promoter replaced by the HNRPA2B1 promoter. Expression and infection of retrovirus performed following standard protocols. Whole-cell extracts from cells expressing empty constructs, ERG-FLAG or EHF-FLAG were run on SDS-PAGE gels and blotted to nitrocellulose membranes following standard procedures. Antibodies used for immunodetection were FLAG (M2, Sigma Life Sciences) and beta-actin (C4, Fisher Scientific).

### Chromatin Immunoprecipitation (ChIP) and ChIPseq analysis

ChIPs were performed as described previously (53), with the following modifications. Cross-linked chromatin was sheared with a Branson sonifier and magnetic beads were washed with buffer containing 500 mM LiCl. Antibodies used for ChIP were: anti-Flag (M2, Sigma Life Sciences) and anti-cJUN (E254, Abcam). ChIPseq libraries were prepared using the NEBNext^®^ ChIP-Seq Library Prep Master Mix Set for Illumina (NEB, E6240) and run on a Hiseq2000 sequencer. Sequence reads were aligned with Novoalign to human genome HG19 and enriched regions (peaks) determined using the MACS2 analysis package (54). Heatmaps of enriched regions for ERG-FLAG and EHF-FLAG ChIPseq were generated with DeepTools (55) using a bedfile corresponding to coordinates from the combined ChIPseq bedfiles, and bigwig files generated from the individual ChIPseq datasets. Data were aligned using the center point of this shared peak bedfile.

Overrepresented DNA sequences present in the FLAG-ERG and FLAG-EHF enriched regions were determined using the MEME-ChIP program (56) (http://meme-suite.org) using default settings except for following parameters for MEME: 1) any number of repetitions for site distribution; and 2) maximum site width of 13. ETS-AP1 sites spacings were determined using Regulatory Sequence Analysis Tools (57) searching the top 1000 ChIPseq peaks for the FLAG-ERG and FLAG-EHF datasets with the string indicated in Figure 3B.

Primer-BLAST (58) was used to generate primer sets for amplification of enriched regions; primer sequences and the coordinates of interrogated regions are provided in Table S9. qPCR of ChIP DNA was performed using Roche FastStart Essential DNA Green Master and run on a Roche Lightcycler 96. Serially diluted input was used to create a standard curve for absolute quantitation of amplified regions from ChIP DNA. PCRs for each sample and primer pair were run as triplicates and signal averaged over the three values. Data are displayed in graphical form as a ratio of the signal of the target region over the signal of a negative control genomic region. An input sample was also subject to same qPCR reactions graphed to confirm validity of negative control region. For all primer pairs, the input enrichment value was approximately one.

### Structural modeling

Structural models for ETS and JUN-FOS factors binding to a composite DNA sequence were constructed with PyMOL (Version 1.7.0.5) and the following PDB entries: ERG, 4IRI (25); JUN-FOS, 1FOS (24); ELF3, 3JTG (26). A homology model of EHF was generated from the closely related ELF3 by manual mutation of distinct residues. ETS and JUN-FOS molecules were oriented by aligning DNA nucleotides to create a ETS-AP1 composite sequence, such as those found in the *UPP* promoter and the *COPS8* enhancer (Table S8).

### Protein Atlas and cBioPortal data curation

Protein levels for ETS factors in normal prostate and prostate cancer samples were curated from The Protein Atlas (https://www.proteinatlas.org)(27,28). Data are reported for ELF1, ELF2, ELF4, and SPIB; these ETS factors, or close homologs (Figs. S5, S7), have been shown to bind to ETS-AP1 DNA in an anticooperative manner with JUN-FOS here or previously (17). These four factors are expressed at a “medium” level in normal prostate cells so “low” or “no detection” in prostate cancer cells represents downregulation at the protein level. TCGA data were curated from cBioPortal (http://www.cbioportal.org) (29,30). Several prostate cancer genomic studies revealed recurrent gene deletions of ELF1 in up to 20% of patient samples (31–36). An example from one study is represented in Fig. 6C and all studies with substantial ELF1 deletions are listed in Fig. S8B.

## Acknowledgements

We thank Mahesh Chandrasekharan, Jedediah Doane, and members of the Graves lab for helpful discussion. This work was supported by the National Institutes of Health (R01GM38663 to B.J.G. and R01CA204121 to P.C.H.). Support to B.J.G. from the Huntsman Cancer Institute/Huntsman Cancer Foundation and the Howard Hughes Medical Institute is acknowledged. Shared resources at the University of Utah were supported by the National Institutes of Health (P30CA042014 to the Huntsman Cancer Institute). The content of this publication is solely the responsibility of the authors and does not necessarily represent the official views of the National Institutes of Health or other funding agencies.

## Author Contributions

P.C.H., B.J.G., and S.L.C. conceived the project. B.J.M., K.A.C., N.B., and S.L.C. designed and conducted experiments. All authors analyzed the data. K.A.C., B.J.G., and S.L.C. wrote the manuscript and all authors helped revise.

The authors declare that they have no conflicts of interest with the contents of this article.

## References

1. Lee, T. I., and Young, R. A. (2000) Transcription of eukaryotic protein-coding genes. Annu Rev Genet 34, 77–137

2. Panne, D. (2008) The enhanceosome. Curr Opin Struct Biol 18, 236–242

3. Garvie, C. W., Pufall, M. A., Graves, B. J., and Wolberger, C. (2002) Structural analysis of the autoinhibition of Ets-1 and its role in protein partnerships. J Biol Chem 277, 45529–45536

4. Jolma, A., Yin, Y., Nitta, K. R., Dave, K., Popov, A., Taipale, M., Enge, M., Kivioja, T., Morgunova, E., and Taipale, J. (2015) DNA-dependent formation of transcription factor pairs alters their binding specificity. Nature 527, 384–388

5. LaRonde-LeBlanc, N. A., and Wolberger, C. (2003) Structure of HoxA9 and Pbx1 bound to DNA: Hox hexapeptide and DNA recognition anterior to posterior. Genes Dev 17, 2060–2072

6. Shrivastava, T., Mino, K., Babayeva, N. D., Baranovskaya, O. I., Rizzino, A., and Tahirov, T. H. (2014) Structural basis of Ets1 activation by Runx1. Leukemia 28, 2040–2048

7. Joshi, R., Passner, J. M., Rohs, R., Jain, R., Sosinsky, A., Crickmore, M. A., Jacob, V., Aggarwal, A. K., Honig, B., and Mann, R. S. (2007) Functional specificity of a Hox protein mediated by the recognition of minor groove structure. Cell 131, 530–543

8. Kim, S., Brostromer, E., Xing, D., Jin, J., Chong, S., Ge, H., Wang, S., Gu, C., Yang, L., Gao, Y. Q., Su, X. D., Sun, Y., and Xie, X. S. (2013) Probing allostery through DNA. Science 339, 816–819

9. Panne, D., Maniatis, T., and Harrison, S. C. (2007) An atomic model of the interferon-beta enhanceosome. Cell 129, 1111–1123

10. Shiina, M., Hamada, K., Inoue-Bungo, T., Shimamura, M., Uchiyama, A., Baba, S., Sato, K., Yamamoto, M., and Ogata, K. (2015) A novel allosteric mechanism on protein-DNA interactions underlying the phosphorylation-dependent regulation of Ets1 target gene expressions. J Mol Biol 427, 1655–1669

11. Hollenhorst, P. C., Ferris, M. W., Hull, M. A., Chae, H., Kim, S., and Graves, B. J. (2011) Oncogenic ETS proteins mimic activated RAS/MAPK signaling in prostate cells. Genes Dev 25, 2147–2157

12. Bosc, D. G., Goueli, B. S., and Janknecht, R. (2001) HER2/Neu-mediated activation of the ETS transcription factor ER81 and its target gene MMP-1. Oncogene 20, 6215–6224

13. Tugores, A., Le, J., Sorokina, I., Snijders, A. J., Duyao, M., Reddy, P. S., Carlee, L., Ronshaugen, M., Mushegian, A., Watanaskul, T., Chu, S., Buckler, A., Emtage, S., and McCormick, M. K. (2001) The epithelium-specific ETS protein EHF/ESE-3 is a context-dependent transcriptional repressor downstream of MAPK signaling cascades. J Biol Chem 276, 20397–20406

14. Plotnik, J. P., Budka, J. A., Ferris, M. W., and Hollenhorst, P. C. (2014) ETS1 is a genome-wide effector of RAS/ERK signaling in epithelial cells. Nucleic Acids Res 42, 11928–11940

15. Tomlins, S. A., Rhodes, D. R., Perner, S., Dhanasekaran, S. M., Mehra, R., Sun, X. W., Varambally, S., Cao, X., Tchinda, J., Kuefer, R., Lee, C., Montie, J. E., Shah, R. B., Pienta, K. J., Rubin, M. A., and Chinnaiyan, A. M. (2005) Recurrent fusion of TMPRSS2 and ETS transcription factor genes in prostate cancer. Science 310, 644–648

16. Tomlins, S. A., Mehra, R., Rhodes, D. R., Smith, L. R., Roulston, D., Helgeson, B. E., Cao, X., Wei, J. T., Rubin, M. A., Shah, R. B., and Chinnaiyan, A. M. (2006) TMPRSS2:ETV4 gene fusions define a third molecular subtype of prostate cancer. Cancer Res 66, 3396–3400

17. Kim, S., Denny, C. T., and Wisdom, R. (2006) Cooperative DNA binding with AP-1 proteins is required for transformation by EWS-Ets fusion proteins. Mol Cell Biol 26, 2467–2478

18. Albino, D., Longoni, N., Curti, L., Mello-Grand, M., Pinton, S., Civenni, G., Thalmann, G., D’Ambrosio, G., Sarti, M., Sessa, F., Chiorino, G., Catapano, C. V., and Carbone, G. M. (2012) ESE3/EHF controls epithelial cell differentiation and its loss leads to prostate tumors with mesenchymal and stem-like features. Cancer Res 72, 2889–2900

19. Cangemi, R., Mensah, A., Albertini, V., Jain, A., Mello-Grand, M., Chiorino, G., Catapano, C. V., and Carbone, G. M. (2008) Reduced expression and tumor suppressor function of the ETS transcription factor ESE-3 in prostate cancer. Oncogene 27, 2877–2885

20. Cheng, X. H., Black, M., Ustiyan, V., Le, T., Fulford, L., Sridharan, A., Medvedovic, M., Kalinichenko, V. V., Whitsett, J. A., and Kalin, T. V. (2014) SPDEF inhibits prostate carcinogenesis by disrupting a positive feedback loop in regulation of the Foxm1 oncogene. PLoS Genet 10, e1004656

21. Nye, J. A., Petersen, J. M., Gunther, C. V., Jonsen, M. D., and Graves, B. J. (1992) Interaction of murine ets-1 with GGA-binding sites establishes the ETS domain as a new DNA-binding motif. Genes Dev 6, 975–990

22. Wei, G. H., Badis, G., Berger, M. F., Kivioja, T., Palin, K., Enge, M., Bonke, M., Jolma, A., Varjosalo, M., Gehrke, A. R., Yan, J., Talukder, S., Turunen, M., Taipale, M., Stunnenberg, H. G., Ukkonen, E., Hughes, T. R., Bulyk, M. L., and Taipale, J. (2010) Genome-wide analysis of ETS-family DNA-binding in vitro and in vivo. EMBO J 29, 2147–2160

23. Hollenhorst, P. C., Chandler, K. J., Poulsen, R. L., Johnson, W. E., Speck, N. A., and Graves, B. J. (2009) DNA specificity determinants associate with distinct transcription factor functions. PLoS Genet 5, e1000778

24. Glover, J. N., and Harrison, S. C. (1995) Crystal structure of the heterodimeric bZIP transcription factor c-Fos-c-Jun bound to DNA. Nature 373, 257–261

25. Regan, M. C., Horanyi, P. S., Pryor, E. E., Jr., Sarver, J. L., Cafiso, D. S., and Bushweller, J. H. (2013) Structural and dynamic studies of the transcription factor ERG reveal DNA binding is allosterically autoinhibited. Proc Natl Acad Sci U S A 110, 13374–13379

26. Agarkar, V. B., Babayeva, N. D., Wilder, P. J., Rizzino, A., and Tahirov, T. H. (2010) Crystal structure of mouse Elf3 C-terminal DNA-binding domain in complex with type II TGF-beta receptor promoter DNA. J Mol Biol 397, 278–289

27. Uhlen, M., Zhang, C., Lee, S., Sjostedt, E., Fagerberg, L., Bidkhori, G., Benfeitas, R., Arif, M., Liu, Z., Edfors, F., Sanli, K., von Feilitzen, K., Oksvold, P., Lundberg, E., Hober, S., Nilsson, P., Mattsson, J., Schwenk, J. M., Brunnstrom, H., Glimelius, B., Sjoblom, T., Edqvist, P. H., Djureinovic, D., Micke, P., Lindskog, C., Mardinoglu, A., and Ponten, F. (2017) A pathology atlas of the human cancer transcriptome. Science 357

28. Uhlen, M., Fagerberg, L., Hallstrom, B. M., Lindskog, C., Oksvold, P., Mardinoglu, A., Sivertsson, A., Kampf, C., Sjostedt, E., Asplund, A., Olsson, I., Edlund, K., Lundberg, E., Navani, S., Szigyarto, C. A., Odeberg, J., Djureinovic, D., Takanen, J. O., Hober, S., Alm, T., Edqvist, P. H., Berling, H., Tegel, H., Mulder, J., Rockberg, J., Nilsson, P., Schwenk, J. M., Hamsten, M., von Feilitzen, K., Forsberg, M., Persson, L., Johansson, F., Zwahlen, M., von Heijne, G., Nielsen, J., and Ponten, F. (2015) Proteomics. Tissue-based map of the human proteome. Science 347, 1260419

29. Cerami, E., Gao, J., Dogrusoz, U., Gross, B. E., Sumer, S. O., Aksoy, B. A., Jacobsen, A., Byrne, C. J., Heuer, M. L., Larsson, E., Antipin, Y., Reva, B., Goldberg, A. P., Sander, C., and Schultz, N. (2012) The cBio cancer genomics portal: an open platform for exploring multidimensional cancer genomics data. Cancer Discov 2, 401–404

30. Gao, J., Aksoy, B. A., Dogrusoz, U., Dresdner, G., Gross, B., Sumer, S. O., Sun, Y., Jacobsen, A., Sinha, R., Larsson, E., Cerami, E., Sander, C., and Schultz, N. (2013) Integrative analysis of complex cancer genomics and clinical profiles using the cBioPortal. Sci Signal 6, pl1

31. Robinson, D., Van Allen, E. M., Wu, Y. M., Schultz, N., Lonigro, R. J., Mosquera, J. M., Montgomery, B., Taplin, M. E., Pritchard, C. C., Attard, G., Beltran, H., Abida, W., Bradley, R. K., Vinson, J., Cao, X., Vats, P., Kunju, L. P., Hussain, M., Feng, F. Y., Tomlins, S. A., Cooney, K. A., Smith, D. C., Brennan, C., Siddiqui, J., Mehra, R., Chen, Y., Rathkopf, D. E., Morris, M. J., Solomon, S. B., Durack, J. C., Reuter, V. E., Gopalan, A., Gao, J., Loda, M., Lis, R. T., Bowden, M., Balk, S. P., Gaviola, G., Sougnez, C., Gupta, M., Yu, E. Y., Mostaghel, E. A., Cheng, H. H., Mulcahy, H., True, L. D., Plymate, S. R., Dvinge, H., Ferraldeschi, R., Flohr, P., Miranda, S., Zafeiriou, Z., Tunariu, N., Mateo, J., Perez-Lopez, R., Demichelis, F., Robinson, B. D., Sboner, A., Schiffman, M., Nanus, D. M., Tagawa, S. T., Sigaras, A., Eng, K. W., Elemento, O., Sboner, A., Heath, E. I., Scher, H. I., Pienta, K. J., Kantoff, P., de Bono, J. S., Rubin, M. A., Nelson, P. S., Garraway, L. A., Sawyers, C. L., and Chinnaiyan, A. M. (2015) Integrative Clinical Genomics of Advanced Prostate Cancer. Cell 162, 454

32. Cancer Genome Atlas Research, N. (2015) The Molecular Taxonomy of Primary Prostate Cancer. Cell 163, 1011–1025

33. Grasso, C. S., Wu, Y. M., Robinson, D. R., Cao, X., Dhanasekaran, S. M., Khan, A. P., Quist, M. J., Jing, X., Lonigro, R. J., Brenner, J. C., Asangani, I. A., Ateeq, B., Chun, S. Y., Siddiqui, J., Sam, L., Anstett, M., Mehra, R., Prensner, J. R., Palanisamy, N., Ryslik, G. A., Vandin, F., Raphael, B. J., Kunju, L. P., Rhodes, D. R., Pienta, K. J., Chinnaiyan, A. M., and Tomlins, S. A. (2012) The mutational landscape of lethal castration-resistant prostate cancer. Nature 487, 239–243

34. Baca, S. C., Prandi, D., Lawrence, M. S., Mosquera, J. M., Romanel, A., Drier, Y., Park, K., Kitabayashi, N., MacDonald, T. Y., Ghandi, M., Van Allen, E., Kryukov, G. V., Sboner, A., Theurillat, J. P., Soong, T. D., Nickerson, E., Auclair, D., Tewari, A., Beltran, H., Onofrio, R. C., Boysen, G., Guiducci, C., Barbieri, C. E., Cibulskis, K., Sivachenko, A., Carter, S. L., Saksena, G., Voet, D., Ramos, A. H., Winckler, W., Cipicchio, M., Ardlie, K., Kantoff, P. W., Berger, M. F., Gabriel, S. B., Golub, T. R., Meyerson, M., Lander, E. S., Elemento, O., Getz, G., Demichelis, F., Rubin, M. A., and Garraway, L. A. (2013) Punctuated evolution of prostate cancer genomes. Cell 153, 666–677

35. Kumar, A., Coleman, I., Morrissey, C., Zhang, X., True, L. D., Gulati, R., Etzioni, R., Bolouri, H., Montgomery, B., White, T., Lucas, J. M., Brown, L. G., Dumpit, R. F., DeSarkar, N., Higano, C., Yu, E. Y., Coleman, R., Schultz, N., Fang, M., Lange, P. H., Shendure, J., Vessella, R. L., and Nelson, P. S. (2016) Substantial interindividual and limited intraindividual genomic diversity among tumors from men with metastatic prostate cancer. Nat Med 22, 369–378

36. Taylor, B. S., Schultz, N., Hieronymus, H., Gopalan, A., Xiao, Y., Carver, B. S., Arora, V. K., Kaushik, P., Cerami, E., Reva, B., Antipin, Y., Mitsiades, N., Landers, T., Dolgalev, I., Major, J. E., Wilson, M., Socci, N. D., Lash, A. E., Heguy, A., Eastham, J. A., Scher, H. I., Reuter, V. E., Scardino, P. T., Sander, C., Sawyers, C. L., and Gerald, W. L. (2010) Integrative genomic profiling of human prostate cancer. Cancer Cell 18, 11–22

37. Hollenhorst, P. C., McIntosh, L. P., and Graves, B. J. (2011) Genomic and biochemical insights into the specificity of ETS transcription factors. Annu Rev Biochem 80, 437–471

38. Hollenhorst, P. C., Jones, D. A., and Graves, B. J. (2004) Expression profiles frame the promoter specificity dilemma of the ETS family of transcription factors. Nucleic Acids Res 32, 5693–5702

39. Bose, R., Karthaus, W. R., Armenia, J., Abida, W., Iaquinta, P. J., Zhang, Z., Wongvipat, J., Wasmuth, E. V., Shah, N., Sullivan, P. S., Doran, M. G., Wang, P., Patruno, A., Zhao, Y., International, S. U. C. P. C. F. P. C. D. T., Zheng, D., Schultz, N., and Sawyers, C. L. (2017) ERF mutations reveal a balance of ETS factors controlling prostate oncogenesis. Nature 546, 671–675

40. Sgouras, D. N., Athanasiou, M. A., Beal, G. J., Jr., Fisher, R. J., Blair, D. G., and Mavrothalassitis, G. J. (1995) ERF: an ETS domain protein with strong transcriptional repressor activity, can suppress ets-associated tumorigenesis and is regulated by phosphorylation during cell cycle and mitogenic stimulation. EMBO J 14, 4781–4793

41. Stephens, D. N., Klein, R. H., Salmans, M. L., Gordon, W., Ho, H., and Andersen, B. (2013) The Ets transcription factor EHF as a regulator of cornea epithelial cell identity. J Biol Chem 288, 34304–34324

42. Ando, M., Kawazu, M., Ueno, T., Koinuma, D., Ando, K., Koya, J., Kataoka, K., Yasuda, T., Yamaguchi, H., Fukumura, K., Yamato, A., Soda, M., Sai, E., Yamashita, Y., Asakage, T., Miyazaki, Y., Kurokawa, M., Miyazono, K., Nimer, S. D., Yamasoba, T., and Mano, H. (2016) Mutational Landscape and Antiproliferative Functions of ELF Transcription Factors in Human Cancer. Cancer Res 76, 1814–1824

43. Sizemore, G. M., Pitarresi, J. R., Balakrishnan, S., and Ostrowski, M. C. (2017) The ETS family of oncogenic transcription factors in solid tumours. Nat Rev Cancer 17, 337–351

44. Goetz, T. L., Gu, T. L., Speck, N. A., and Graves, B. J. (2000) Auto-inhibition of Ets-1 is counteracted by DNA binding cooperativity with core-binding factor alpha2. Mol Cell Biol 20, 81–90

45. Shah, N. H., Wang, Q., Yan, Q., Karandur, D., Kadlecek, T. A., Fallahee, I. R., Russ, W. P., Ranganathan, R., Weiss, A., and Kuriyan, J. (2016) An electrostatic selection mechanism controls sequential kinase signaling downstream of the T cell receptor. Elife 5

46. Shah, N. H., Lobel, M., Weiss, A., and Kuriyan, J. (2018) Fine-tuning of substrate preferences of the Src-family kinase Lck revealed through a high-throughput specificity screen. Elife 7

47. Selvaraj, N., Kedage, V., and Hollenhorst, P. C. (2015) Comparison of MAPK specificity across the ETS transcription factor family identifies a high-affinity ERK interaction required for ERG function in prostate cells. Cell Commun Signal 13, 12

48. Currie, S. L., Lau, D. K. W., Doane, J. J., Whitby, F. G., Okon, M., McIntosh, L. P., and Graves, B. J. (2017) Structured and disordered regions cooperatively mediate DNA-binding autoinhibition of ETS factors ETV1, ETV4 and ETV5. Nucleic Acids Res 45, 2223–2241

49. Studier, F. W. (2005) Protein production by auto-induction in high density shaking cultures. Protein Expr Purif 41, 207–234

50. Currie, S. L., Doane, J. J., Evans, K. S., Bhachech, N., Madison, B. J., Lau, D. K. W., McIntosh, L. P., Skalicky, J. J., Clark, K. A., and Graves, B. J. (2017) ETV4 and AP1 Transcription Factors Form Multivalent Interactions with three Sites on the MED25 Activator-Interacting Domain. J Mol Biol 429, 2975–2995

51. Ferguson, H. A., and Goodrich, J. A. (2001) Expression and purification of recombinant human c-Fos/c-Jun that is highly active in DNA binding and transcriptional activation in vitro. Nucleic Acids Res 29, E98

52. Benjamini, Y., Krieger, A. M., and Yekutieli, D. (2006) Adaptive linear step-up procedures that control the false discovery rate. Biometrika 93, 491–507

53. Hollenhorst, P. C., Shah, A. A., Hopkins, C., and Graves, B. J. (2007) Genome-wide analyses reveal properties of redundant and specific promoter occupancy within the ETS gene family. Genes Dev 21, 1882–1894

54. Zhang, Y., Liu, T., Meyer, C. A., Eeckhoute, J., Johnson, D. S., Bernstein, B. E., Nusbaum, C., Myers, R. M., Brown, M., Li, W., and Liu, X. S. (2008) Model-based analysis of ChIP-Seq (MACS). Genome Biol 9, R137

55. Ramirez, F., Ryan, D. P., Gruning, B., Bhardwaj, V., Kilpert, F., Richter, A. S., Heyne, S., Dundar, F., and Manke, T. (2016) deepTools2: a next generation web server for deep-sequencing data analysis. Nucleic Acids Res 44, W160–165

56. Machanick, P., and Bailey, T. L. (2011) MEME-ChIP: motif analysis of large DNA datasets. Bioinformatics 27, 1696–1697

57. Thomas-Chollier, M., Sand, O., Turatsinze, J. V., Janky, R., Defrance, M., Vervisch, E., Brohee, S., and van Helden, J. (2008) RSAT: regulatory sequence analysis tools. Nucleic Acids Res 36, W119–127

58. Ye, J., Coulouris, G., Zaretskaya, I., Cutcutache, I., Rozen, S., and Madden, T. L. (2012) Primer-BLAST: a tool to design target-specific primers for polymerase chain reaction. BMC Bioinformatics 13, 134

